# Chronic intermittent ethanol produces nociception through endocannabinoid-independent mechanisms in mice

**DOI:** 10.1101/2024.11.08.622656

**Authors:** C Miliano, Y Dong, M Proffit, N Corvalan, LA Natividad, AM Gregus, MW Buczynski

## Abstract

Alcohol use disorder (AUD) affects millions of people and represents a significant health and economic burden. Pain represents a frequently under-treated aspect of hyperkatifeia during alcohol withdrawal, yet to date no drugs have received FDA approval for the treatment of this indication in AUD patients. This study aims to evaluate the potential of targeting bioactive lipid signaling pathways as a therapeutic approach for treating alcohol withdrawal-related pain. We utilized a chronic intermittent ethanol (CIE) vapor exposure model in C57BL/6J mice of both sexes to establish alcohol dependence, and demonstrated that CIE mice developed robust tactile allodynia and thermal hyperalgesia during withdrawal that was independent of prior blood alcohol levels. Next, we evaluated four drugs for their efficacy in reversing tactile allodynia during abstinence from CIE using a cross-over treatment design that included FDA-approved naltrexone as well as commercially available inhibitors targeting inflammatory lipid signaling enzymes including fatty acid amide hydrolase (FAAH), monoacylglycerol lipase (MAGL), and 15-Lipoxygenase (LOX). None of these compounds produced significant therapeutic benefit in reversing established CIE-induced tactile allodynia, despite attenuating pain-like behaviors at these doses in other chronic pain models. Additionally, we assessed plasma endocannabinoid levels in both sexes during withdrawal. We found that there is an inherent sex difference in the endogenous anti-inflammatory endocannabinoid tone in naive mice and CIE treatment affected endocannabinoids levels in female mice only. These findings underscore the need to better understand the driving causes of AUD induced pain and to develop novel therapeutic approaches to mitigate pain in AUD patients.

## Introduction

Alcohol use disorder (AUD) is a persistent public health issue that impacts approximately 400 million people worldwide, including 28.9 million individuals in the US in 2023^1^. AUD is characterized by a cycle of binge/intoxication, withdrawal/negative affect, and preoccupation/anticipation^1^ that perpetuates problematic alcohol use, with the withdrawal/negative affect aspect of this cycle being a powerful driver behind relapse and continued use^2^. Among the patients who were formally diagnosed and sought treatment, 33% experienced additional AUD episodes even after they had been asymptomatic for 12 months or more^3^, indicating a necessity for novel therapeutics to treat alcohol use disorder.

Pain is an under-treated aspect of alcohol withdrawal that motivates relapse and continued use. Due in part to the acute antinociceptive effects of alcohol on pain, the misuse of alcohol in an attempt to manage pain as well as disruptions to daily life due to pain are prevalent in problem drinkers^4^, with chronic pain serving as a predictor for relapse^5^. AUD patients are more likely to seek care to address health concerns like chronic pain that result from long-term AUD than excessive alcohol use^6,7^, and a significant portion of treatment-seeking patients with AUD experience recurring pain^8^. Chronic alcohol consumption can result in peripheral neuropathy^9^ and alcohol withdrawal directly leads to an increase in pain hypersensitivity^10^. Preclinical rodent models can recapitulate these symptoms, as chronic intermittent ethanol vapor exposure (CIE) model of ethanol exposure and withdrawal reliably produces increased nociception in both rats and mice, similar to the experience of human AUD patients^11,12^. While there is a clear emerging connection between pain and problematic alcohol use, none of the current FDA approved medications for AUD mitigate pain experienced by patients^13–15^, and other off-label pain medications often cannot be used long-term^16,17^ or carry substantial risk of producing additional dependence^18,19^.

Endocannabinoids and related bioactive lipids have emerged as potential targets for treating AUD withdrawal-induced chronic pain by altering inflammatory processes. Patients report that self-medication with cannabinoid-based products may reduce alcohol intake or even promote abstinence^20,21^ as well as minimize use of opioid analgesics^22,23^. However, the use of cannabinoid-based treatments poses additional concerns of dependence for an already vulnerable population^24,25^. In contrast, elevation of endogenous cannabinoids represents an alternative therapeutic approach that could attenuate AUD-induced chronic pain while circumventing safety concerns due to dependence. Increases of endogenous anandamide (AEA) has been shown to reduce AUD-induced anxiety-like behaviors and alcohol intake in rat and mouse models of alcohol dependence^26^ while increases of endogenous 2-arachidonoylglycerol (2-AG) has similar therapeutic benefits on anxiety-like behaviors and alcohol intake in rodent models of alcohol dependence^26^. Another N-acyl ethanolamide - Oleoylethanolamide (OEA) - has been shown to regulate multiple aspects of alcohol exposure in preclinical studies^27,28^ and both OEA and Palmitoylethanolamide (PEA) have been reported to be elevated in binge drinking patients^27^. Previous studies in humans also suggest that circulating bioactive lipids such as endocannabinoids and eicosanoids are altered during abstinence and may predict craving in humans^29,30^. Inhibition of fatty acid amide hydrolase (FAAH) and monoacylglycerol lipase (MAGL), key enzymes regulating endogenous endocannabinoid levels, have been shown to reduce inflammatory^31–33^ and neuropathic^34,35^ pain-like behaviors in rodents via elevating levels of AEA and 2-AG, and FAAH inhibitors have reversed pain-like behaviors in mice subjected to the two-bottle choice (2BC) model of chronic ethanol exposure^36^. Furthermore, we have demonstrated that in AUD patients, plasma levels of pro-nociceptive 12/15-Lipoxygenase (LOX) metabolites predict alcohol craving during abstinence^30^. However, therapeutics targeting these lipid pathways have not been evaluated for any potential anti-nociceptive effects in a preclinical model of chronic alcohol consumption-induced dependence.

Importantly, despite the significant prevalence of AUD among women, there is a lack of knowledge on sex differences in pharmacotherapies for AUD^37,38^. Women tend to experience more severe health consequences from alcohol consumption compared to men such as liver damage^39^, heart disease^40^ and brain damage^41^. Additionally, women have not been included in most of the AUD clinical trials^37^. Therefore, preclinical studies including both sexes are critical to underline potential mechanisms unique to women.

The present study aims to assess the potential for targeting bioactive lipid signaling pathways as a therapeutic approach for treating alcohol withdrawal-related pain in both male and female mice. We established conditions for a reliable mouse model of alcohol dependence with robust and recurring expression of tactile and thermal pain-like behaviors during withdrawal using the chronic intermittent ethanol (CIE) vapor exposure model in mice of both sexes. Using this approach, FDA-approved naltrexone as well as commercially available inhibitors targeting FAAH, MAGL, or 15-LOX were evaluated as potential treatments for pain hypersensitivity during withdrawal using a cross-over treatment design. Lastly, blood samples were collected at the end of the experimental procedures to assess plasma levels of AEA, 2-AG, OEA, and PEA as well as other bioactive lipids belonging to the eicosanoids class.

## Methods

### Animals

C57BL/6J male and female mice (10 weeks old with an average body weight of 26g and 20g respectively at initiation of the experiment) were obtained from Jackson Labs. Mice were housed in individually ventilated cages under sanitary conditions at 3-5 mice per cage, in a 12h reverse dark-light cycle (22:00 on/10:00 off), temperature and humidity-controlled room. Mice have access to food and water *ad libitum*. All protocols and experiments were approved by the Virginia Tech Institutional Animal Care and Use Committee (IACUC) and complied with the ARRIVE guidelines^42^.

### Drugs

Naltrexone hydrochloride (Sigma-Aldrich N3136, Source BCCF6115) was dissolved in saline (0.9% Sodium Chloride Injection, USP, ICU Medical Inc. Cat. # 798437, NDC 0990-7983-02). ML351 (15-LOX-1 inhibitorMedChemExpress Cat. #HY-111310, Lot # 110818), PF-3845 (FAAH inhibitor, Cayman Chemical Company Cat. # 13279, Batch: 0454664-38, 0454664-50), or MJN110 (MGL inhibitor, MedChemExpress Cat. # HY-117474, lot # 150639) were dissolved in DMSO (Sigma, Cat. # 41640), mixed with Tween-80 (Sigma, Cat. # P9416) at a 1:1 ratio and then diluted in water and followed by PBS for a final ratio of 1:1:9:9 (DMSO : Tween80 : Water : PBS, or 5% DMSO/5%Tween-20). All lipid primary and internal standards were purchased from Cayman Chemicals. Formic acid was purchased from Sigma Aldrich. All HPLC-grade solvents (Water, Acetonitrile, Ethanol, Isopropyl alcohol) were purchased from VWR.

### Chronic Ethanol Exposure (CIE) Model

Mice were assigned to either ethanol vapor group (CIE) or air group (AIR) based on body weight, tactile withdrawal thresholds at baseline, and hot plate response time at baseline to avoid confounding effects. Alcohol exposure was performed following the established chronic ethanol vapor exposure (CIE) paradigm (Fig 1) as previously published^11,12,43^. Briefly, mice were exposed to vaporized ethanol or air for 16 hours overnight using a vacuum-based vapor exposure system (La Jolla Alcohol Research) and returned to housing cages for 8 hours, for a total of four days per week (one week is considered one vaping cycle). To produce ethanol vapor, an HPLC pump (Waters, 515 HPLC Pump) was used to supply 95% ethanol (Pharmco, ethyl alcohol 95%, 190 proof; Cat. # 111000190) into a vaporizer (Glas-Col, heating mantle, Cat. #: 100B TM106) connected to the vacuum-based system. Mice of the CIE group were injected intraperitoneally (IP, 10 ml/kg) with alcohol (1.75 g/kg, in 0.9% saline solution) and pyrazole (alcohol dehydrogenase inhibitor, Sigma-Aldrich P56607, Source BCCK9523; 68.1 mg/kg, in 0.9% saline solution) before being placed (by cages of 5) into ethanol exposure chambers. Mice of the AIR group were injected with saline (0.9% solution) and pyrazole (68.1 mg/kg) before being placed into air exposure chambers. For all mice, cheek blood was collected immediately after the first and third 16-hour exposure to ethanol vapor or air in each vaping cycle to determine blood alcohol levels (BAL, mg/dL). The flowrate of ethanol was adjusted to maintain a target BAL of 150 – 225 mg/dL during cycles of CIE exposure^12,44^. Mice were weighed daily prior to the start of vaping sessions during each vaping cycle.

**Figure 1.**
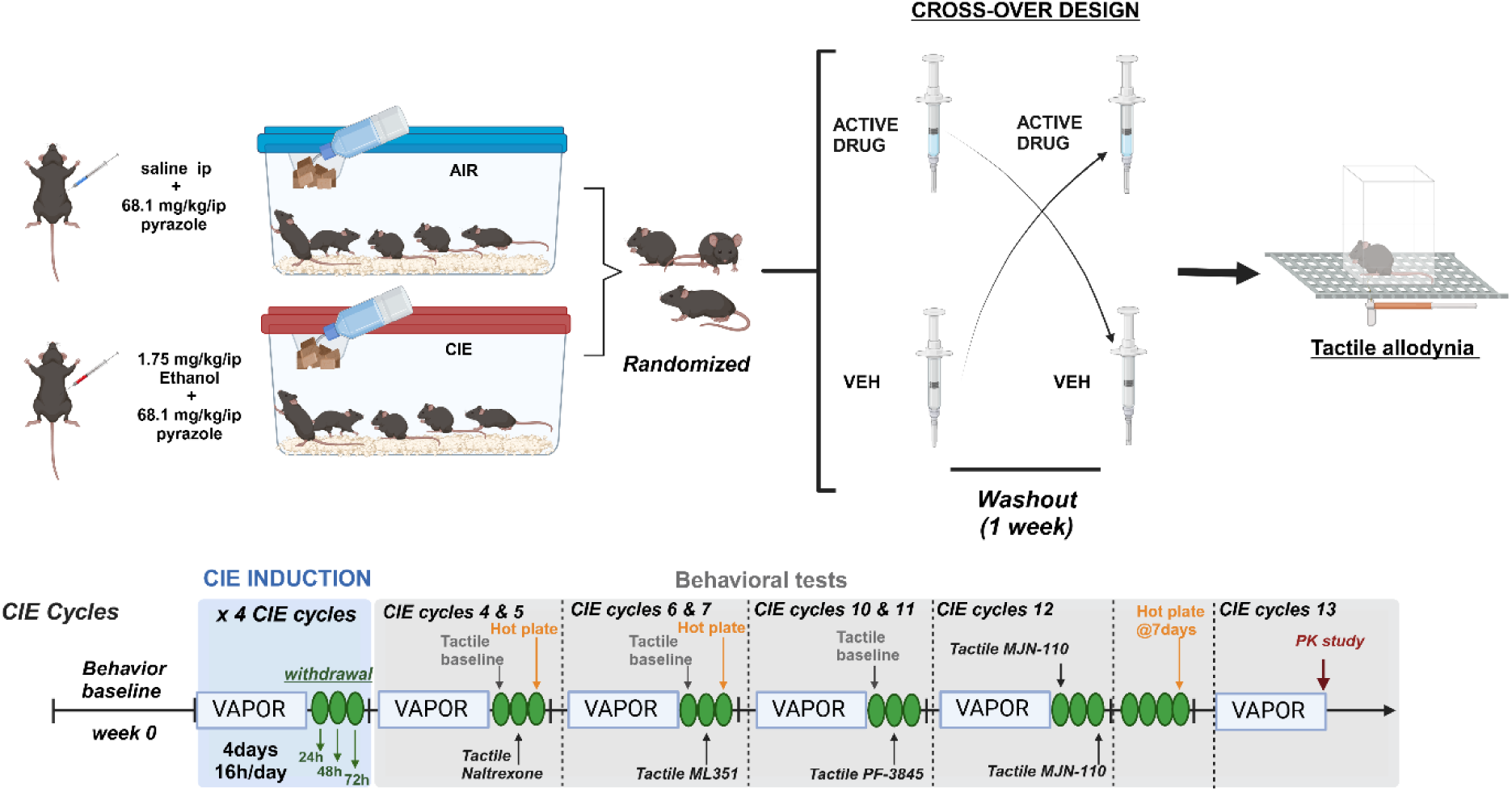
Representative CIE Timeline and experimental design. Alcohol exposure was performed following the established Chronic Intermittent Ethanol vapor exposure model where mice were exposed to either room air control (AIR) or vaporized ethanol (CIE) for 12 to13 cycles. Each cycle consisted of 4 consecutive days of exposure (16 hours overnight) with preexposure administration of pyrazole (68.1 mg/kg) and alcohol (1.75 g/kg, IP), followed by 3 days of abstinence. Therapeutic treatments were performed using a crossover design where all mice received either vehicle of drug treatment on separate testing days. This figure was created using BioRender.com. IP, intraperitoneal.

### Blood Ethanol Level by Gas Chromatography

Levels of ethanol in whole blood samples were assessed using GC-FID with a method adapted from Gonzalez *et. al.*^45^ and Doyon *et. al*^46^. For all CIE and AIR mice, 10 μL of blood was collected and mixed with 90uL of saturated sodium chloride solution (technical, ∼26%; Sigma-Aldrich, Cat. # 71392) in a 2 mL glass mass spectrometry sample vial (2mL Clear Glass 12x32mm Flat Base 9-425 Screw Thread Vial; ALWSCI technologies, Cat. # C0000008). Following collection, the sample was capped and heated at 50°C for 20 minutes, before manual sampling using Solid Phase MicroExtraction (SPME) fiber (d_f_ 75 μm, carboxen/polydimethylsiloxane; Supelco, Cat. # 57318) for 15 seconds. Blood ethanol level (mg/dL) was determined using a HP 6890 GC-FID with an Agilent DB-ALC2 column (30m × 0.320mm × 1.20μm) as the stationary phase and Helium as the mobile phase. ChemStation software (v A.10.02) was used to analyze the resulting ethanol peaks. External standards were assessed for calibration each time (quantitative curve between 25 and 400 mg/dL).

### Thermal Escape Latency (hot plate test)

Thermal hyperalgesia was assessed using the hot plate test as previously published^47^. Briefly, mice received room habituation for an hour with cage lids off prior to individual apparatus habituation, where each mouse was placed within an acrylic chamber (IITC, Part #39ME, 9 cm diameter, 30 cm height) on top of the hot plate apparatus (IITC, Part #39) for 10 minutes while it is turned off. Room lighting was maintained between 35 - 40 lux. Thermal escape latency (time in seconds to shaking, lifting, or licking hind paws, and jumping) was recorded using a hand timer for each animal. Baseline measurement was recorded prior to alcohol exposure to determine the paw withdrawal latency (PWL) of each mouse. Recordings were conducted at temperatures of 50°C, 52.5°C, and 55°C with a cutoff time of 20 seconds to prevent tissue damage. Each mouse was tested three times at each temperature on separate days, with each test taking place at 72h after the last alcohol exposure. The average of three measures was taken as the mean thermal PWL of the mouse at a given temperature.

### Tactile Allodynia (von Frey test)

Tactile allodynia was assessed with the up-down method utilizing manual von Frey filaments as previously published^47^. Briefly, filaments with buckling forces ranging from 0.02 and 2 g were used to assess the average 50% paw withdrawal threshold (PWT) of each hindpaw at baseline and at timepoints throughout the experiment as indicated. Mice were habituated to the testing room and chambers for 2 h for up to 5 days prior to baseline and then 1 h on test day. Any mouse with a baseline 50% PWT ≤ 0.79 g was excluded from the study. As alcohol exposure was systemic, PWT for both hindpaws were averaged, and data was presented using the 50% gram threshold vs time, or area under the curve (AUC). Additionally, the % maximum possible effect (MPE) was calculated for each animal over the course of testing, and the difference between CIE % MPE and AIR % MPE average (delta % MPE) for each animal was calculated, both as previously described^48^.

#### Measurement of Endocannabinoids and Bioactive Lipids in Plasma Samples

Submandibular blood samples were collected as previously described^49^ from mice of both sexes and groups 24 hours after the last exposure, processed into plasma, and stored at −80 °C until liquid chromatography mass spectrometry lipid analysis was performed as previously published ^30^ and described in Supplemental Information.

### Statistical Analysis

All statistical analyses were performed using GraphPad Prism (v 10.3.1) and detailed reports are available in Supplemental Tables 1–3. Data are presented as mean ± SEM, with discrete data points included where appropriate. For the pharmacokinetic (PK) study, a non-linear one-phase decay fit was used. Behavioral test assessments, including baseline and experimental conditions, were analyzed using two-way ANOVA (tactile pre-test thresholds factors: weeks × CIE exposure; hot plate factors: sex × CIE exposure at each temperature/withdrawal duration), followed by Tukey’s (tactile pre-test thresholds) and Bonferroni’s (hot plate) post hoc test. Pearson’s R correlation was used for correlation studies. Unpaired t-test was used to assess the effect of sex on delta % MPE of tactile thresholds. Mixed effects analysis followed by Bonferroni’s post-hoc test was applied for the tactile crossover study within each sex and treatment condition (factors: drug treatment × CIE exposure). Plasma lipid levels are reported as pg/ml and were analyzed by using two-way ordinary ANOVA (factors: sex × CIE exposure) followed by Tukey’s post hoc test. Statistical significance was defined as *P < 0.05, **P < 0.01, ***P < 0.001, and ****P < 0.0001.

## Results

### Chronic intermittent ethanol vapor exposure (CIE) pharmacokinetics

In this study, we used a chronic intermittent ethanol (CIE) vapor paradigm of alcohol dependence in C57BL/6J male and female mice (Fig 1) as previously described^43^. Mice were exposed to up to 13 cycles of CIE, and compared with control mice exposed to air without alcohol (AIR) throughout the study. The CIE mice displayed average BALs between 150-225 mg/dl at each CIE cycle starting at week 2 to 4 (Fig 2A,B), a paradigm previously shown to establish alcohol dependence in mice^12,43^. In order to produce dependence, the CIE model incorporates daily pyrazole administration which can influence BAL and alcohol-induced pain-like responding in mice ^11,12,43^. Thus, we conducted a post-vapor exposure BAL time-course to evaluate the impact of pyrazole on ethanol pharmacokinetics in CIE mice of both sexes. In males, following a vapor exposure session (Fig 2,D), CIE mice (T_1/2_ = 2.0 h) metabolize alcohol similar to naïve mice (Fig 2C) in the absence of pyrazole (T_1/2_ = 1.2 h) and faster than naïve mice immediately after pyrazole treatment (T_1/2_ = 3.4 h). In females instead, CIE mice metabolize alcohol in a slower way (Fig 2D) (T_1/2_ = 5.0 h) compared to naïve mice (Fig 2C) in the absence of pyrazole (T_1/2_ = 1.4 h) and naïve mice immediately after pyrazole treatment (T_1/2_ = 3.6 h). Collectively, these data confirm that CIE mice exhibit near complete metabolism of alcohol between vaping sessions, and indicate that nociceptive behaviors emergent during withdrawal can be evaluated without the confounding anti-nociceptive influence of alcohol starting at 24 h of abstinence.

**Figure 2.**
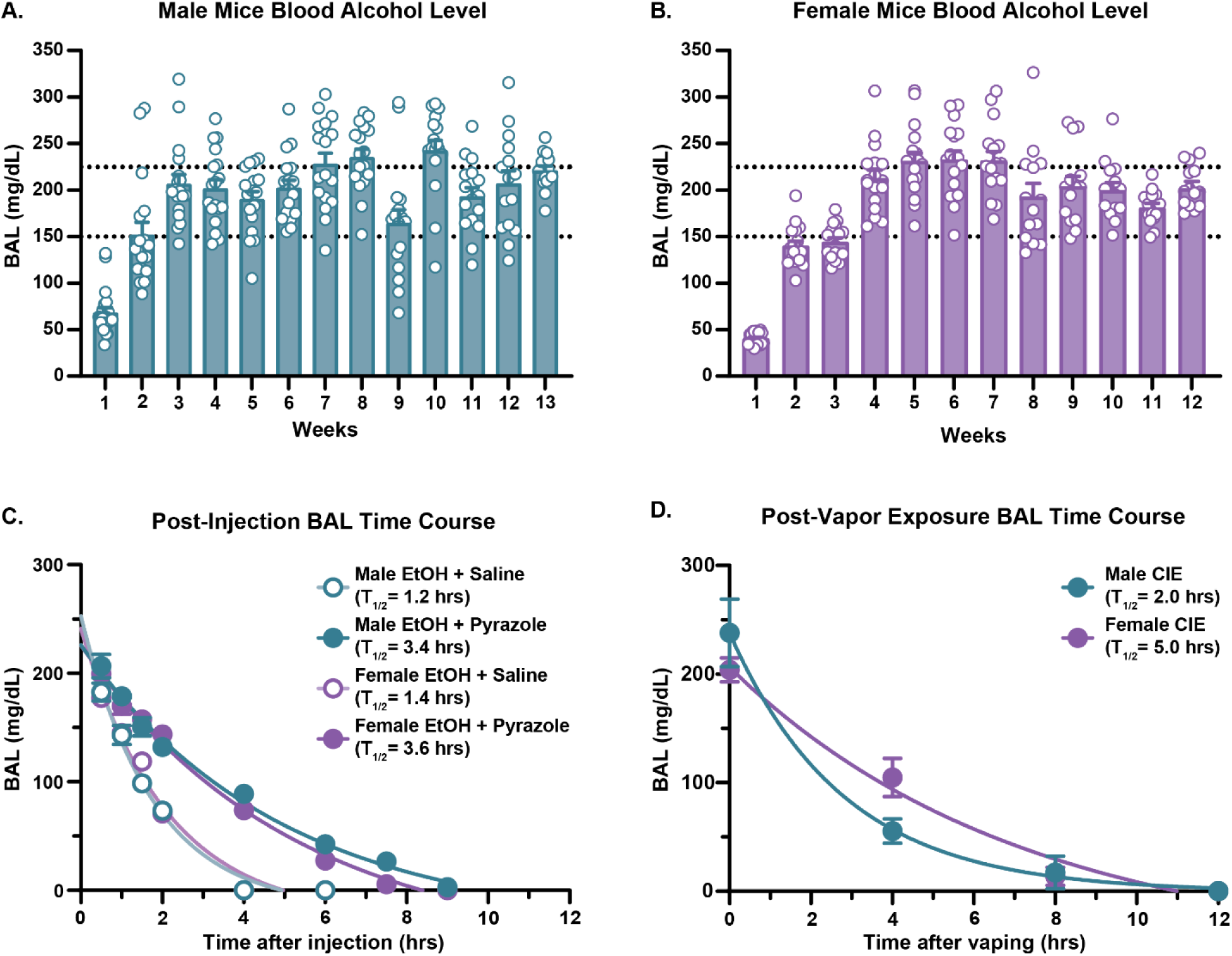
Establishment of the CIE paradigm. (A-B) BAL time course of mice exposed to vaporized ethanol for 16 hours overnight following an IP injection of ethanol and pyrazole; **(A)** male mice (n = 15-20 mice/group), **(B)** female mice (n = 14-20 mice/group). **(C)** Time Course following IP EtOH injection paired with saline or IP EtOH paired with pyrazole for determination of half-life (male: n = 5-6 mice/group, female: n = 4-5 mice/group). **(D)** Time Course and respective half-life following overnight 16-hour ethanol vaping session paired with IP injection of EtOH and pyrazole (male: n = 4-5 mice/group, female: n = 5 mice/group). Data presented as mean ± SEM. Significance indicated by *P < 0.05, **P < 0.01. CIE, Chronic Intermittent Ethanol vapor exposure; BAL, Blood Alcohol Level; IP, intraperitoneal; EtOH, ethanol.

### CIE induces tactile and thermal pain hypersensitivity during abstinence in male and female mice

Next, we validated that mice develop robust and persistent tactile allodynia during abstinence following at least 4 cycles of CIE^11,12^. To this end, we measured baseline tactile paw withdrawal thresholds (PWT) as depicted in Fig 3A prior to initiating the CIE vapor exposure paradigm and subsequently tested mice 24h after the end of each cycle to determine alcohol withdrawal-induced changes to PWT. We found a significant sex difference between the delta % MPE of tactile thresholds as a result of CIE withdrawal, with a greater effect of withdrawal on the tactile thresholds of female mice (Fig 3B, Table S2). We also identified a significant effect of group and time whereby CIE withdrawal precipitated a reduction in tactile thresholds starting at the third cycle for males and 4th cycle for females, achieving maximal effect from the 4th cycle until the end of the study (Fig 3C, D, Table S2). Since alcohol itself exhibits acute analgesic effects, we hypothesized that the immediately prior BAL may correlate with the subsequent pain levels that emerge during withdrawal. For this reason, we performed a Pearson correlation test between each mouse’s tactile threshold during abstinence with the prior BAL during CIE (Fig 3E, F, Table S2). We found no significant correlation between the tactile allodynia during abstinence and their prior BAL, suggesting that the tactile allodynia observed during withdrawal is independent of the acute analgesic effects of alcohol.

**Figure 3.**
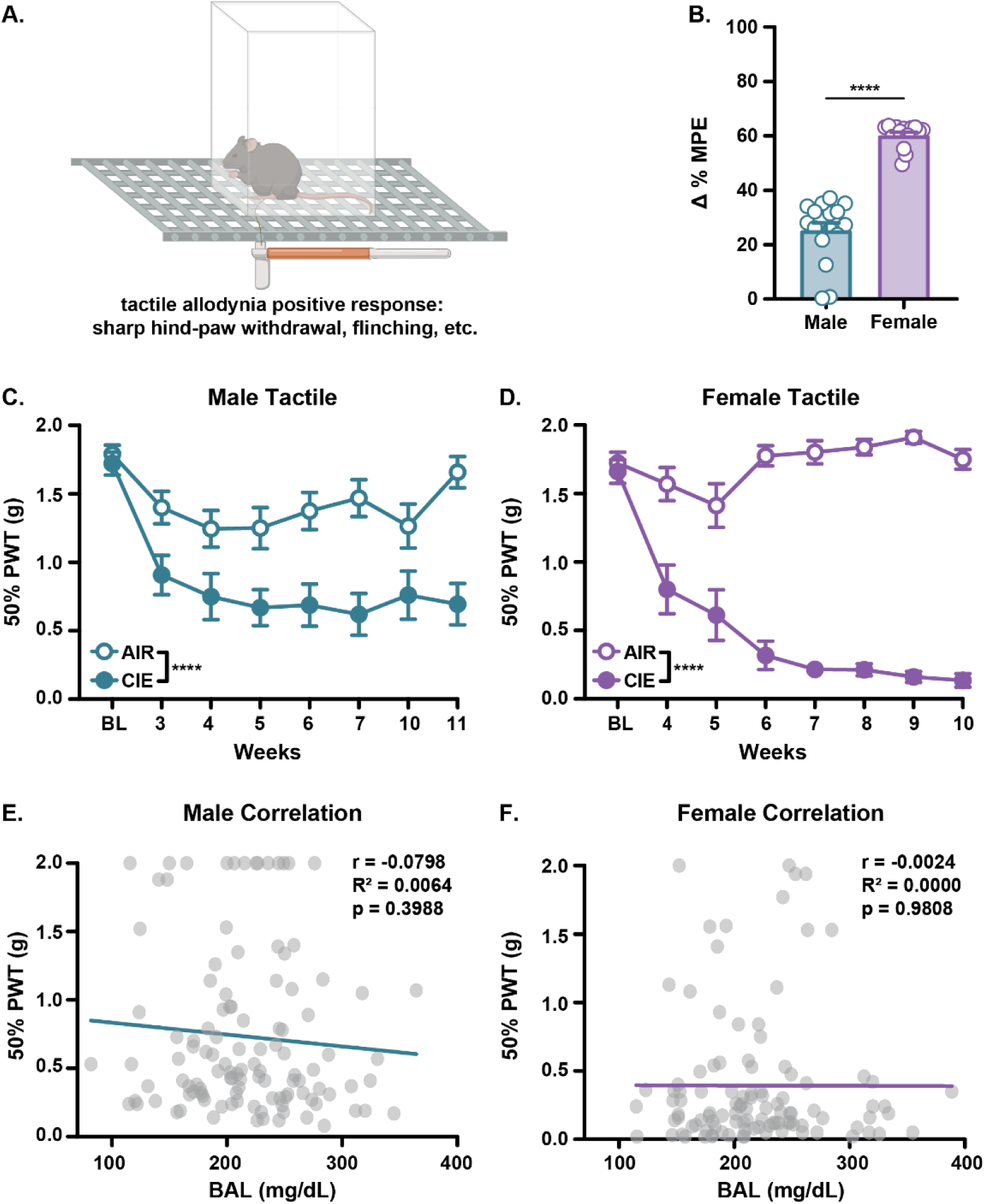
Measurement of tactile allodynia during abstinence using the CIE paradigm. **(A)** Illustration of the tactile test setup. **(B)** change in tactile threshold % MPE due to alcohol exposure in male and female mice, calculated from tactile timecourses for **(C)** male (n = 15-20 mice/group) and **(D)** female (n = 14-20 mice/group) mice. Correlation graphs comparing individual mouse BAL to its corresponding PWT result for **(E)** male mice (n = 15-18 mice/group) and **(F)** female (n = 14-17 mice/group) mice. Data presented as mean ± SEM. Significance indicated by **P<0.01, ****P<0.0001. n = 15-20 mice/group. MPE, maximum possible effect; CIE, Chronic Intermittent Ethanol vapor exposure; AUC, Area Under the Curve; PWT, Paw Withdrawal Threshold; BAL, Blood Alcohol Level.

Previous studies reported mixed findings regarding the effect of alcohol exposure on thermal hyperalgesia in mice^12^. These discrepancies may result from the thermal stimulus intensity required to unmask a hyperalgesic response during alcohol withdrawal, and thus we performed an assessment of thermal paw withdrawal latencies (PWL) using the hot plate at three different calibrated temperatures (50°C, 52.5°C, 55°C) as depicted in Fig 4A. At 55°C, we identified expression of thermal hyperalgesia in both male and female CIE mice at 72h of withdrawal that persisted for at least 7 days (Fig 4B, Table S2). There was also a main effect of sex with female mice showing a significantly lower duration overall compared to the male animals at both time points (Fig 4B, Table S2). Interestingly, while this main effect of sex persisted at all experimental conditions (Fig 4B – D, Table S2), only female mice demonstrated thermal hyperalgesia at 52.5°C at 72h which persisted for at least 7 days, and there was no expression of thermal hyperalgesia at this temperature for the male mice (Fig 4C, Table S2). Neither sex showed expression of thermal hyperalgesia at 50°C in agreement with previous reports^12^ (Fig 4D, Table S2). To confirm that prior single day BALs do not strongly influence the subsequent expression of thermal hyperalgesia that emerges during withdrawal, we performed a Pearson correlation between each mouse’s thermal response latency (at 55°C) during abstinence with the prior day BAL during CIE and found no significant effect (Fig. 4E, F). Collectively, these data demonstrate the development of post-CIE tactile and thermal pain hypersensitivity in both sexes of mice using this paradigm.

**Figure 4.**
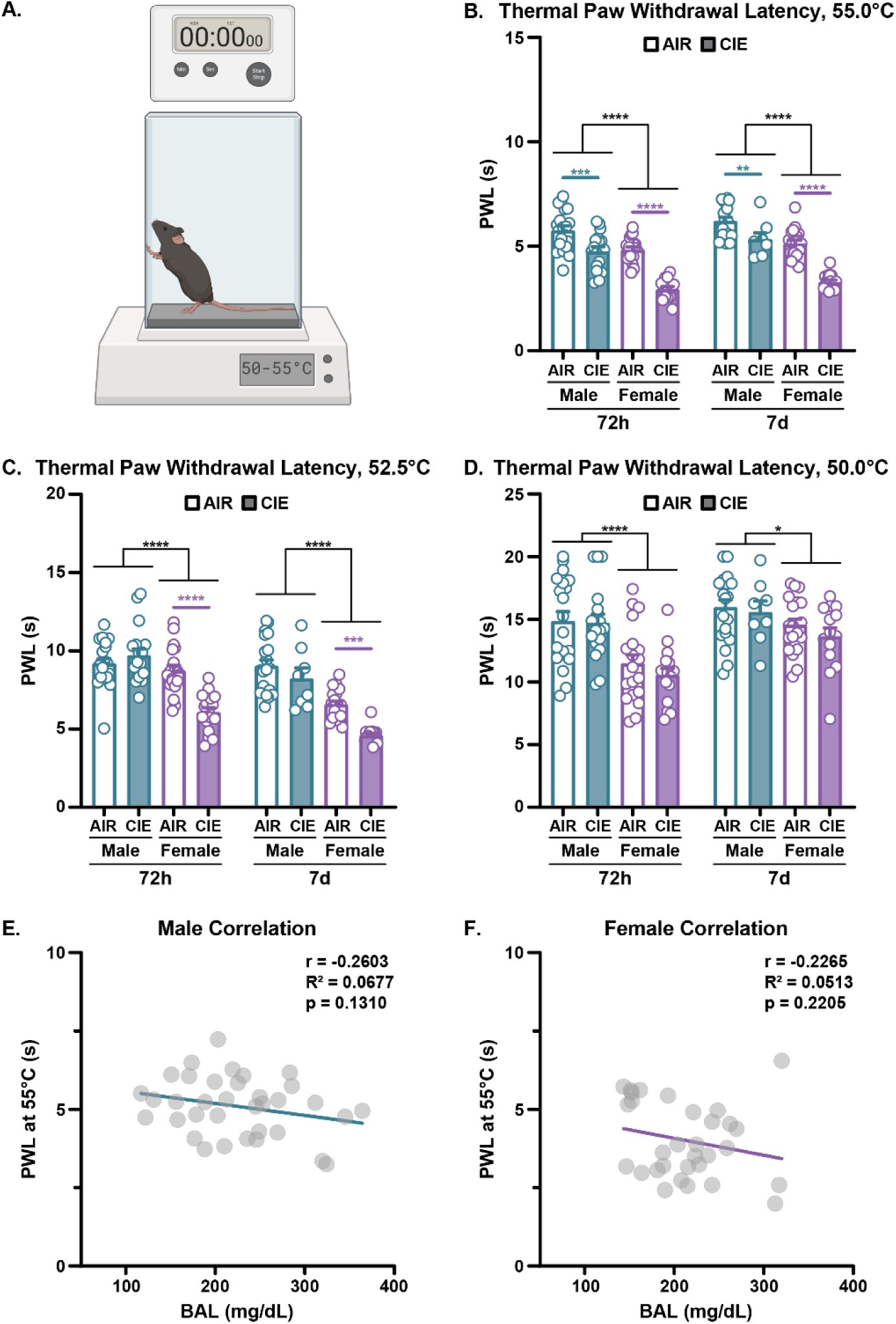
Measurement of thermal hyperalgesia during abstinence using the CIE paradigm. **(A)** Illustration of the hot plate test setup. **(B-D)** Thermal Paw Withdrawal Latencies of male and female mice, unexposed (AIR) or exposed (CIE) to alcohol, assessed at various temperatures at 72h and 7 days following last exposure at **(B)** 55°C, **(C)** 52.5°C, or **(D)** 50°C. **(E)** Correlation graph comparing individual male mice BAL to its corresponding thermal PWL result (n = 17-18 mice/group). **(F)** Correlation graph comparing individual female mice BAL to its corresponding thermal PWL result (n = 14-16 mice/group). Significance indicated by *P < 0.05, **P < 0.0, ** P < 0.001, ****P < 0.0001. CIE, Chronic Intermittent Ethanol vapor exposure; PWL, Paw Withdrawal Latency; BAL, Blood Alcohol Level.

### Targeting pro-inflammatory signaling did not reverse CIE tactile allodynia

While currently there are 3 FDA-approved drugs for treating AUD, none of these has been shown to attenuate hyperkatifeia by reversing withdrawal-induced pain hypersensitivity. Using our established chronic vapor AUD model, we conducted a series of 4 cross-over therapeutic studies (Fig 5, Table S2) where both AIR and CIE mice received either vehicle or drug treatment on alternating testing cycles at 48h from the last alcohol vapor exposure. First, we evaluated the efficacy of naltrexone in reducing hyperkatifeia because it is an FDA-approved treatment for AUD, reduces ethanol intake in both preclinical AUD models and clinical studies of AUD patients^37,50–54^, and has been suggested as a potential therapeutic approach for treating inflammation and chronic pain^55^. Following systemic treatment with naltrexone at a dose that effectively reduces alcohol consumption (3 mg/kg, IP, 2h prior to testing)^50,51^, tactile PWT were not significantly different in AIR- or CIE-treated male (Fig 5A, Table S2) or female (Fig. 5E, Table S2) mice, indicating no effect of opioid receptor antagonism on hyperkatifeia in this model.

**Figure 5.**
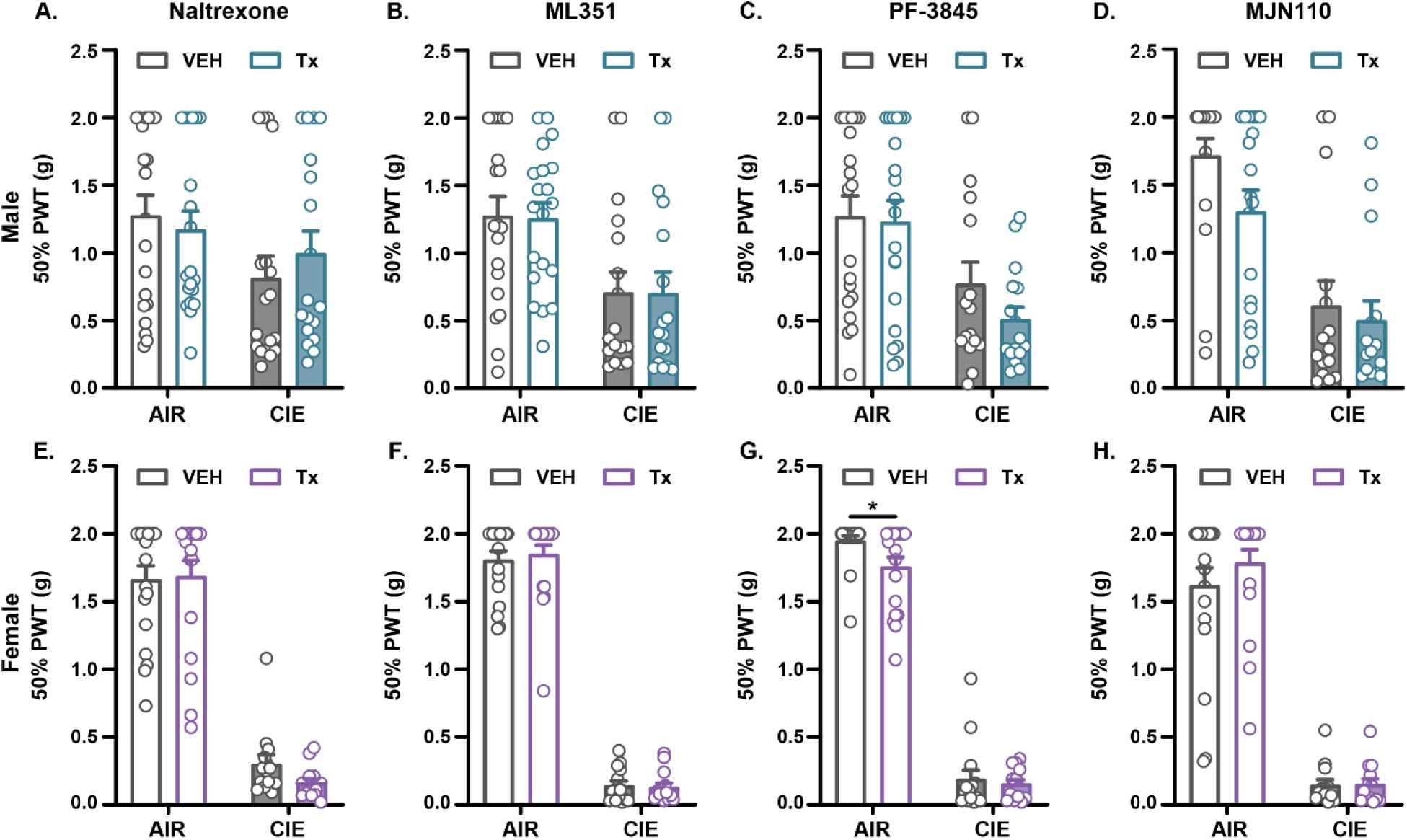
Anti-hyperalgesic drugs targeting inflammatory signaling do not reverse CIE-induced tactile allodynia. 50% Tactile Paw Withdrawal Thresholds for vehicle v. drug-treated **(A-D)** male mice (AIR, n = 19 - 20; CIE, n = 15 - 18) or **(E-H)** female mice (AIR, n = 19; CIE, n = 14) unexposed (AIR) or exposed (CIE) to alcohol. **(A, E)** Naltrexone: 3 mg/kg IP; **(B,F)** ML351: 30 mg/kg IP; **(C, G)** PF-3845: 10 mg/kg IP; **(D, H)** MJN110: 10 mg/kg IP. Data presented as mean ± SEM. Significance indicated by *P < 0.05. PWT, CIE, Chronic Intermittent Ethanol vapor exposure; Paw Withdrawal Threshold; VEH, vehicle; Tx, Drug Treatment; IP, intraperitoneal.

Second, we chose to examine the effects of a selective inhibitor of 15-LOX (ML351) on hyperkatifeia. 15-LOX synthesizes pro-inflammatory lipid metabolites including 15-HETE which we found to be associated with alcohol craving in humans^30^. Additionally, we have shown that 15-LOX inhibition reverses tactile allodynia in a preclinical model of neuropathic pain-like behaviors in rats and mice^56,57^. Following systemic treatment with an anti-hyperalgesic dose of ML351 (30 mg/kg, IP, 1h prior to testing), tactile PWT were not significantly different in AIR- or CIE-treated male (Fig 5B, Table S2) or female (Fig 5F, Table S2) mice, indicating no effect of 15-LOX inhibition on hyperkatifeia in this model. These results show that two treatments with therapeutic benefit in other chronic pain models do not influence the AUD pain state that develops following CIE.

### Targeting anti-inflammatory lipid signaling did not reverse CIE tactile allodynia

Given the established preclinical success in targeting endocannabinoid signaling for treating other aspects of AUD as well as other models of chronic pain, we interrogated these pathways as a potential treatment option for the CIE-induced pain state in mice. Pro-resolving endocannabinoids such as anandamide (AEA) and 2-arachidonoylglycerol (2-AG) may affect AUD-induced chronic pain as these bioactive lipids reduce alcohol intake in rodents^58,59^ and elicit antinociceptive effects in other preclinical models of chronic pain^33,36,60,61^. First, we evaluated the effects of elevating endogenous AEA levels using the selective fatty-acid amide hydrolase (FAAH) inhibitor PF-3845 because it reduces ethanol intake in rodent AUD dependence models^26^, decreases withdrawal signs including anxiety-like and depression-like behaviors^26^, reverses pain-like behaviors in models of inflammatory pain^60,61^, and an optimized clinical candidate derived from this compound successfully advanced through Phase 1 Clinical Trials^62^. Following systemic treatment with a dose of PF-3845 that effectively reduces alcohol consumption (10 mg/kg, IP, 2h prior to testing), tactile PWT were unchanged in both AIR- and CIE-treated male mice (Fig 5C, Table S2). Similarly, PWT were unchanged in CIE-treated yet were reduced in AIR control female mice (Fig 5G, Table S2), indicating no specific effect of FAAH inhibition on hyperkatifeia in this model.

Finally, we examined the effects of elevating endogenous 2-AG levels using the selective monoacylglycerol lipase inhibitor (MAGL) MJN110 as it also reduces ethanol intake in rodent AUD dependence models^26,63^, decreases withdrawal signs including anxiety-like and depression-like behaviors^26,63^, reverses pain-like behaviors during alcohol withdrawal^36,64^ and during chemotherapy-induced peripheral neuropathy^35^, and a humanized MAGL inhibitor has advanced successfully through Phase 1 Clinical Trials^65,66^. Following systemic treatment with MJN110 at a dose that effectively reduces alcohol consumption (10 mg/kg, IP, 2h prior to testing), tactile PWT were not significantly different in AIR- or CIE-treated male (Fig 5D, Table S2) or female (Fig 5H, Table S2) mice. Collectively, these findings suggest that elevating endogenous anti-inflammatory endocannabinoids with FAAH or MAGL inhibitors does not reverse the CIE-induced chronic pain state in mice.

### Quantification of anti-inflammatory lipid levels in the plasma during withdrawal

Given the established role of endocannabinoid signaling in AUD as well as other models of chronic pain, we measured blood levels of these anti-inflammatory lipids (AEA, 2-AG, PEA, and OEA) at a time point at which mice displayed tactile allodynia. Our findings show that while CIE treatment did not affect plasma AEA levels in either sex, there was a significant baseline sex difference, with plasma AEA levels being lower in female v. male AIR groups (Fig 6A, Table S3). Plasma 2-AG showed a significant main effect of sex as levels in both AIR and CIE groups of female mice were higher than in their respective counterpart treatment groups of male mice (Fig 6B, Table S3). Plasma PEA levels displayed a significant main effect of sex as levels in both AIR and CIE groups of female mice were lower than in their respective counterpart treatment groups of male mice, while CIE treatment increased PEA in females only (Fig 6C, Table S3). Similarly, we observed a sex difference in plasma OEA levels for CIE-treated males v. females as well as a significant effect of CIE treatment in female mice (Fig 6D, Table S3). Collectively, these findings suggest that there are sex differences in the endogenous anti-inflammatory endocannabinoid tone (in females: lower for AEA and PEA and higher for 2-AG and OEA) and that CIE treatment elevates PEA and OEA levels in female mice. The results for other bioactive lipids belonging to the eicosanoid family are reported in Table S3.

**Figure 6.**
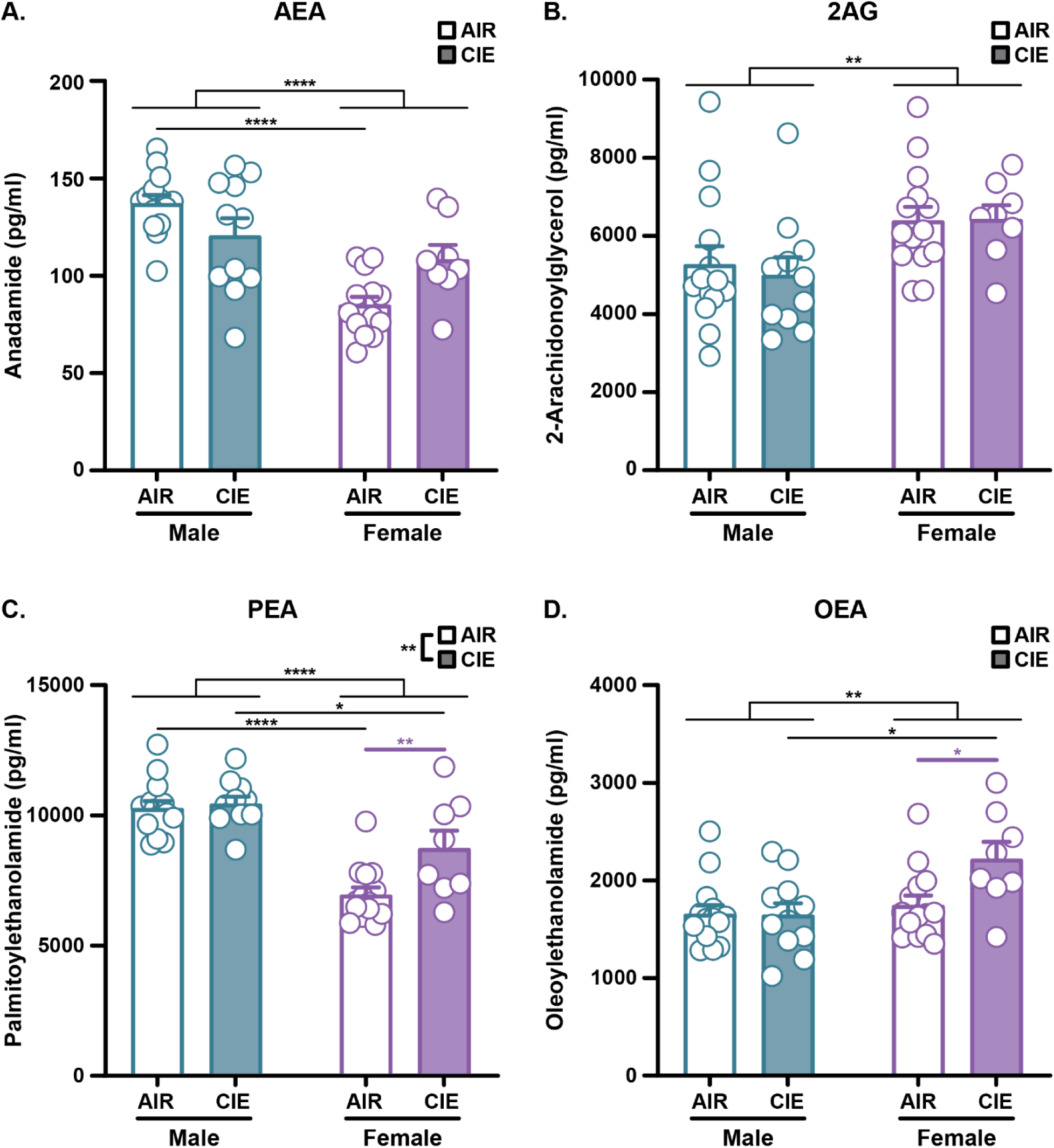
Anti-inflammatory lipid levels in the plasma during withdrawal. **(A)** AEA, **(B)** 2-AG, **(C)** PEA, and **(D)** OEA levels at 24h since the last exposure presented as pg/ml. (n = 14, AIR; n = 8 - 11, CIE). Significance indicated by *P < 0.05, **P<0.01, ****P<0.0001. CIE, Chronic Intermittent Ethanol vapor exposure; AEA, anandamide; 2-AG, 2-arachidonoylglycerol; PEA, Palmitoylethanolamide; OEA, Oleoylethanolamide.

## Discussion

Pain represents an important component of alcohol dependence and withdrawal that motivates relapse and continued use, yet to date no therapeutic options have been identified for addressing this aspect of hyperkatifeia. In this study, we implemented the CIE vapor model of alcohol dependence in male and female C57BL/6J mice, demonstrating that this paradigm elicited both tactile allodynia and thermal hyperalgesia during acute abstinence in both sexes. These withdrawal symptoms emerged at 24 hours, lasted for at least 7 days, and persisted throughout 12-13 CIE cycles. Using this model of AUD-induced hyperkatifeia, we performed a multi-drug crossover treatment study using several compounds targeting either pro-inflammatory or pro-resolving signaling pathways. Despite showing therapeutic effects on other models of chronic pain and other output measures of AUD, these drug treatments did not produce a significant reversal of AUD-induced tactile allodynia in either sex.

While multiple preclinical AUD studies have identified tactile pain hypersensitivity in rodents during abstinence, some evaluating thermal responses in mice report mixed findings. Our study found that CIE led to a small but significant decrease in PWL in mice of both sexes during abstinence using a high intensity thermal stimulus (55°C). This stimulus temperature typically is utilized for examination of analgesic effects of drugs, thereby aligning with published findings that evaluated thermal thresholds in alcohol-exposed mice with the Hargreaves test using similar testing conditions with the glass maintained at 32°C^67^. Alternatively, results from our lab and others^12^ using lower hotplate temperatures (50°C to 52.5°C) unmasked a greater sensitivity in females (at 52.5°C) but no differences at either temperature in males. By contrast, abstinence results in much more robust tactile allodynia across multiple studies despite differing species, sexes, and models of alcohol exposure^11,12,36,64,67–75^. The current study uncovered greater apparent tactile pain hypersensitivity of females during alcohol vapor withdrawal as revealed by the difference in % MPE between AIR controls and CIE. Taken together, these findings support prior claims of a possible role for activation of thinly myelinated A-delta afferent fibers from the dorsal root ganglion in AUD-induced hyperkatifeia as they are sensitive to tactile stimuli, exhibit high temperature thresholds, and are recruited in alcohol-induced neuropathy^76^. In rats, alcohol dependence alters supraspinal circuits responsible for interpretation and response to high intensity thermal stimuli, including exacerbation of pronociceptive central amygdalar projections to the ventrolateral periaqueductal gray^70^ as well as signaling within the lateral habenula^64^. It is important to note that our observations do not preclude the possibility of greater tactile sensitivity of male C57BL/6J mice to CIE occurring at an earlier timepoint (*e.g.* at 4 weeks), or using different forces of von Frey filaments as described previously^12^. Future studies using fully powered groups could be directed toward uncovering mechanistic differences between the sexes during development and maintenance phases of CIE-induced dependence.

Despite demonstration of efficacy in reducing drinking in both preclinical models and clinical studies^37,50–54^, the FDA-approved drug naltrexone did not reverse tactile allodynia following CIE. While low dose naltrexone has been investigated as a potential anti-inflammatory treatment for chronic pain conditions through non-opioid mechanisms^77^, it is not surprising that a higher dosage reflecting treatment for AUD patients would produce a beneficial impact on tactile allodynia as it blocks opioid receptor signaling. In addition, we found that systemic treatment with a 15-LOX inhibitor also did not reverse CIE-induced tactile allodynia in mice. This result is surprising given our previous observations demonstrating a critical role for 15-LOX activity in neuropathic-like pain hypersensitivity^56,57^ and that one of its pro-nociceptive metabolites (15-HETE) predicts craving during abstinence in patients with severe AUD^30^. However, it is possible that hyperalgesia precipitated by alcohol withdrawal differs mechanistically from other chronic pain states and from neural substrates driving craving, as suggested previously^78^.

While there is strong evidence in support of elevating endogenous 2-AG as a strategy for reversing the CIE-induced pain state, acute systemic administration of a therapeutic dose of the selective MAGL inhibitor MJN110 did not attenuate alcohol withdrawal-mediated tactile allodynia. Additionally, plasma levels of 2-AG were unchanged by CIE both in males and females. These observations were surprising as JZL184 (another selective MAGL inhibitor) mitigates other affective behaviors during alcohol withdrawal^26^, and reduced tactile allodynia during abstinence in male mice using the two-bottle choice model of alcohol exposure^36^. In addition, Marchigian Sardinian alcohol-preferring rats of both sexes subjected to alcohol two-bottle choice exhibit tactile allodynia during protracted withdrawal that correlates with decreased 2-AG levels within the lumbar dorsal root ganglion^73^, suggesting a potential for MAGL inhibition to address tactile allodynia that emerges during abstinence. Given that multiple studies indicate that our treatment dose of MJN110 (10 mg/kg, IP) results in near complete inactivation of MAGL^79^ and reduces pain-like behavior in chemotherapy induced neuropathy^35^, the absence of an antihyperalgesic effect is not due to insufficient dosing. It has been suggested that elevated 2-AG signaling may counteract withdrawal-associated tactile allodynia in moderate alcohol exposure models like two-bottle choice, but that alcohol exposure models achieving higher BALs like CIE may override this buffering system^36^. Additionally, other genetic factors may influence the impact of 2-AG on tactile hypersensitivity in AUD, as 2-AG levels in the dorsal root ganglion do not predict tactile allodynia in Wistar rats undergoing alcohol two-bottle choice exposure^73^. It is also possible alcohol leads to circuit-specific changes in different preclinical models of AUD that may not respond to systemic interventions, as bilateral infusions of JZL184 into the lateral habenula (LHb) attenuated CIE-induced pain-like responding Long Evans rats^64^.

It also was somewhat unexpected that treatment with a selective FAAH inhibitor did not reverse tactile allodynia during CIE-induced abstinence even in males, which displayed reduced plasma AEA during withdrawal. Bilateral infusions of the FAAH inhibitor URB597 into the lateral habenula (LHb) attenuated CIE-induced pain-like responding Long Evans rats^64^, and systemic administration of the selective inhibitor PF-3845 reduces alcohol intake and anxiety-like behavior in mice and rats^26^. In the current study, we observed no CIE-induced changes in plasma PEA or OEA in males, and paradoxically increased AEA, PEA and OEA in females, suggesting that anandamide and other ethanolamides may signal through non-cannabinergic pathways during CIE. For example, anandamide can also signal through the pro-nociceptive TRPV1 receptor under certain physiological conditions^80^. Accordingly, alcohol sensitizes TRPV1 in the lateral habenula during alcohol withdrawal to facilitate alcohol-seeking behaviors using the intermittent access two-bottle choice in Long Evans rats^81^. Alternatively, some chronic pain states may be unresponsive to FAAH inhibition. FAAH inhibition did not reverse functional deficits in the monosodium iodoacetate model of osteoarthritis^82^, and a Phase 2 clinical trial using a well-controlled crossover design showed no beneficial effect of FAAH inhibitor PF-04457845 on pain measures despite producing elevated levels of anandamide in these patients. Many of the preclinical pain models where FAAH inhibitors effectively reduce or reverse pain-like behaviors such as carrageenan-induced inflammation^60^ and chronic constriction injury^83^ develop through TLR4-dependent mechanisms^84–86^, and recent work suggests that many symptoms of AUD may develop through TLR4-independent mechanisms^87^. These findings would suggest that alcohol-induced hyperalgesia may represent a distinct mechanism of chronic pain that requires unique therapeutic approaches for treatment.

Taken together, our findings do not support mu opioid receptors, 15-LOX, or endocannabinoid signaling as high value targets for treating hyperalgesia due to severe AUD using the mouse as a model system. However, our results do not preclude the involvement of these mediators in chronic pain experienced by moderate drinkers, nor do they suggest that these pathways do not contribute to other symptoms of AUD. Given the lack of available and effective treatments for AUD-induced pain, future studies should explore new therapeutic approaches using the CIE model.

## Acknowledgements

This work was supported by NIH grants NIAMS R01 AR075241 (AMG); NIDA R00 DA035865 and NCI R01 CA284075 (MWB);

